# Multi temporal multispectral UAV remote sensing allows for yield assessment across European wheat varieties already in tillering stage

**DOI:** 10.1101/2023.05.03.539198

**Authors:** Moritz Camenzind, Kang Yu

**Affiliations:** Precision Agriculture Lab, School of Life Sciences, Technical University of Munich, Freising, Germany; World Agricultural Systems Center (Hans Eisenmann-Forum for Agricultural Sciences – HEF), Technical University of Munich, Freising, Germany

**Keywords:** wheat variety testing, yield prediction, UAV remote sensing, image texture features, machine learning, phenology

## Abstract

High throughput field phenotyping techniques employing multispectral cameras allow to extract a variety of variables and features to predict yield and yield related traits, but little is known about which types of multispectral features may allow to forecast yield potential in the early growth phase. In this study, we hypothesized that the best features for predicting yield in an early stage might be different from the best predictors for the late growth stages. Based on a variety testing trial of 19 European wheat varieties in 2021, multispectral images were taken on 19 dates ranging from tillering to harvest by an unmanned aerial vehicle measuring reflectance in five bands, including visible bands, Red-edge and the near-infrared (NIR). Orthomosaic images were created, and then the single band reflectances, vegetation indices (VI) and texture features (TF) based on a gray level correlation matrix (GLCM) were extracted. We evaluated the performance of these three types of features for yield prediction and classification at different growth stages by, i) using features on each of the measurement dates, ii) smoothing features across the 19 dates, and iii) combining features across the directly adjacent dates, in combination with the random forest models. Our results showed that, for most features, measurements at the flowering stage showed the best performance and the Red reflectance was able to predict yield with a RMSE of 47.4 g m^-2^ (R^2^ = 0.63), the best VI was NDRE predicting yield with a RMSE of 47.9 g m^-2^ (R^2^ = 0.63), the best TF was contrast predicting yield with a RMSE of 57.2 g m^-2^ (R^2^ = 0.46) at the booting stage. Combining dates improved yield prediction in all dates and made the prediction errors more stable across dates. Rather than the Red-edge band, visible bands especially the Red band enabled to distinguish between the high- and low-yielding varieties already in the tillering stage, with a total accuracy of 76.7%. The study confirms our hypothesis and further implies that, in the early stages, the visible bands may be more effective than Red-edge bands in assessing the yield potential in a range of testing varieties.

## 1 Introduction

Improving crop yields in the face of climate change is a significant challenge for plant breeding (Ray et al., 2013). To achieve efficient phenotyping for breeding, it is essential to quickly and accurately identify high-yielding genotypes from a large pool of genotypes (Araus & Cairns, 2014). While traditional phenotyping for variety selection is highly dependent on breeder’s eyes and experiences, high-throughput phenotyping (HTP) emerged in recent years as a more standardizable approach of characterizing plant structure and function and assessing their interactions with the environment by employing various technologies such as imaging, remote sensing, and artificial intelligence (Hund et al., 2019; Watt et al., 2020). HTP-generated high-dimensional phenotypic data embrace the spectral (frequency), spatial, and temporal domains, leading to challenges of analyzing the high dimensional data before they can aid in identifying the genotypes (Bowman et al., 2015; Prey et al., 2022). In particular, field-based HTP including the phenotypic data handling and analysis remain the most significant challenge and requires further development to be fully effective (Kirchgessner et al., 2017; Walter et al., 2015).

Field-based HTP techniques are expected to be fast, cost-effective, and non-destructive (Cabrera-Bosquet et al., 2012). Unmanned aerial systems (UAS-) based remote sensing techniques are increasingly used for HTP of plant traits and yield. Among these sensing techniques, canopy spectral reflectance is highly promising and has been successfully utilized to estimate a diverse range of traits in wheat such as leaf area index (Bukowiecki et al., 2020; Zhang et al., 2021), biomass (Yue et al., 2019), leaf nitrogen and chlorophyll in wheat (Pan et al., 2023) to more general traits such as grain yield and quality (Duan et al., 2017; Prey et al., 2020; Vatter et al., 2022). There are several sensors available for measuring yield under field conditions, such as RGB cameras (Fernandez-Gallego et al., 2019), as well as thermal sensors (Elsayed et al., 2017) and active sensors such as LiDAR (Li et al., 2022). The use of multispectral cameras mounted on unmanned aerial systems (UAS) has proven to be a practical, easy-to-use, and cost-effective approach (Araus et al., 2022).

Multispectral images allow for extracting a variety of features that can be used to predict yield in wheat. Generally, these features can be grouped three categories. Firstly, single band reflectance in specific wavelengths can be directly extracted from multispectral data. Secondly, the reflectance of single bands can be combined to calculate vegetation indices (VIs), which are often more sensitive to specific traits and less affected by environmental conditions during measurement (Tucker, 1979). However, both single band reflectance and VIs may suffer from saturation, particularly for closed canopies (Rischbeck et al., 2016). Thirdly, texture features (TFs) can be extracted to describe the distribution of pixels within a region of interest (ROI). Although TFs can be extracted from any reflectance and VI raster, they often perform less effectively than single band reflectance or VIs in predicting yield in wheat (J. Li et al., 2019). Also, TFs are frequently used in combination with VIs to predict plant traits, e.g., leaf area and biomass in wheat (Zhang et al., 2021). While canopy height often follows a clear temporal dynamics, multispectral features show different dynamics, depending on their sensitivity to a given trait or canopy properties. Also, research indicates that TFs depend heavily on the phenological stage and are therefore their dynamics might interesting to be studied (Culbert et al., 2009). Yet, only limited research focuses on the dynamics of different features in phenotyping, especially the TFs. Among the TF algorithms, one of the mostly used is based on the grey level co-occurrence matrices (GLCMs). In order to calculate GLCMs different parameters such as the level of quantization, the size of the moving window, the moving distance and direction (Haralick et al., 1973). They can be calculated on all available images or rasters of individual bands and VIs. Further, TFs are highly dependent on the GSD and the size of the observed object; and therefore on the camera and the flight height. Zhang et al. (2021) found the best performing TFs for yield prediction were based on the RED as well as the NIR bands. In contrast, Zheng et al. (2019) found that GLCM-based TFs were poorly correlated to above ground biomass (AGB) in rice and calculated normalized difference texture indices that showed a higher correlation with AGB. Further, the phenology has a big influence on the relationship between AGB and the normalized difference texture features (S. Li et al., 2019). Despite that combing feature (e.g., TFs, VIs) and models often improve the prediction, the more reliable performance of yield prediction has been reported in only a few growth stages from the booting to early grain filling (Bowman et al., 2015; Vatter et al., 2022).

Accurately predicting yield using canopy multi-/hyperspectral reflectance further requires careful consideration of the phenological stage of the crop. The anthesis stage or the grain filling stage are often identified as the most suitable stages for yield prediction in wheat (Bowman et al., 2015; Duan et al., 2017; Hassan et al., 2019). In contrast, canopy spectral-based yield prediction has been often reported with a lower accuracy (Prey et al., 2022). Multispectral cameras mounted on UAS enable breeders and researchers to assess the aforementioned spectral and texture features at a high temporal frequency and precision. Within a proper time-window, using a time series for yield prediction allows for the extraction of dynamic canopy traits that could potentially be useful for yield prediction. For instance, Pinter et al. (1981) suggested summing measurement dates after heading to improve yield prediction in wheat and barley. Raun et al. (2001) suggested to take two spectral measurements after dormancy. Time series are further often used to extract information for canopy height, and Taniguchi et al. (2022) used a time-series canopy height model to predict several yield-related traits after the heeding stage. During the stem elongation stage in wheat, Kronenberg et al. (2020) used laser scanning to capture the time series of plant height development and identified quantitative trait loci that accounted for the variability in height dynamic.

Collectively, despite these successes, little is known about which types of multispectral features may allow us to forecast yield potential in the early growth phase. Therefore, this study aims (1) to identify the best performing multispectral traits for yield prediction and classification in wheat (2) to research, if yield types can be classified in relatively early stages and finally (3) to investigate, how traits measured at different time points can be combined to predict yield more accurately.

## 2 Methods

### 2.1 Study site and Environmental monitoring

A trial consisting of 19 diverse European winter wheat elite varieties (*Triticum aestivum*). was grown in plots with a size of 10 m x 1.85 m. The plots were placed in a randomized complete block design with four replicates, resulting in 76 plots totally. The trial took place at the research station of the Technical University of Munich in Dürnast, Freising (48.40630° N, 11.69535° E). The soil at this location can be characterized by a homogeneous Cambisol with 20.8 % clay, 61.5 % silt and 16.6 % sand. All plots were fertilized by applying 180 kg N ha^-^ in three equal splits at BBCH 25, 32 and 65. Plant protection was carried out according to local practice. Sowing took place on the 10.11.2020 and all plots were harvested at full maturity on the 03.08.2021. Precipitation during this period was 1020 mm, the average temperature was 8.2 °C. Climate data was collected from a weather station (Station id 5404) and operated by the Climate Data Center of the German Weather Service. The temperature was aggregated to phenologically meaningful growing degree-days (GDD):

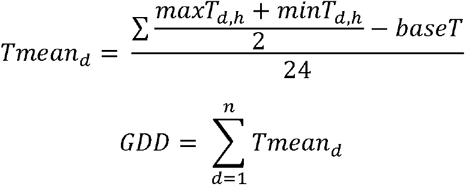

where *Tmean*_*d*_ is the mean temperature for day d after sowing, *maxT*_*d,h*_ and *minT*_*d,h*_ are hourly maximum and minimum temperatures for day d and *baseT* is the base temperature, which was set to 0 °C.

### 2.2 Grain yield, Phenology assessment and Leaf area index measurements

The entire plots were harvested using a combined harvester. The water content of the grains was determined by weighing the grains after harvest, drying them at 65 °C until constant weight was reached and weighing them again. The final yield was normalized to a moisture content of 14 %. The three varieties with the lowest average yield were classified as low yielding and the three varieties with the highest average yield as high yielding. The phenology of each plot was visually rated using the BBCH scale (Meier et al., 2009) on a plot level. Leaf area index (LAI) was measured using a Licor 2000 leaf area meter (LI-COR Biosciences Lincoln, U.S.A.) with a 45° view cap to minimize operator influence. Three measurements were taken at the top of the canopy and four measurements were taken under the canopy at three different locations per plot, which were then averaged.

### 2.3 Multispectral image acquisition and processing

Spectral measurements were aquired using a Phantom 4 Multispectral RTK (DJI, Shenzhen, China). The UAV captures reflectance in wavelengths of 450, 560, 668, 717 and 840 nm and measures the incoming sunlight by a sensor on top of the UAV. Flight height was set to 10 m AGL resulting in a ground sampling distance of 0.7 cm. Overlap in both directions was set to 90 %, the UAV stopped for each image acquisition. Before and after each flight, images of a panel with a known reflectance were taken. Flights were carried out twice per week during heading and flowering stages and once per week at other stages. First flight was carried out on the 25.03.2021 and the last flight on the 20.07.2021, which resulted in 19 flights totally. Images were taken around the solar noon and under sunny conditions, if possible.

The images from each flight were mosaicked using the Agisoft Metashape Professional 1.8.4 (Agisoft, St. Petersburg, Russia) structure-from-motion software. The images were radiometrically calibrated using the reflectance panels on the ground and the incident light sensor on the UAV, with a uniform set of processing parameters used for all flight dates (Figure 1). The point cloud was georeferenced using the real-time kinetic global positioning system (RTK-GPS) integrated into the UAS, with the RTK correction signal provided by SAPOS (Deutsche Landesvermessung). Reflectance of individual bands was extracted by calculating the median of a specific region of interest (ROI) representing a plot using a custom Python 3.7 script (Python Software Foundation, https://www.python.org/).

**Figure 1:**
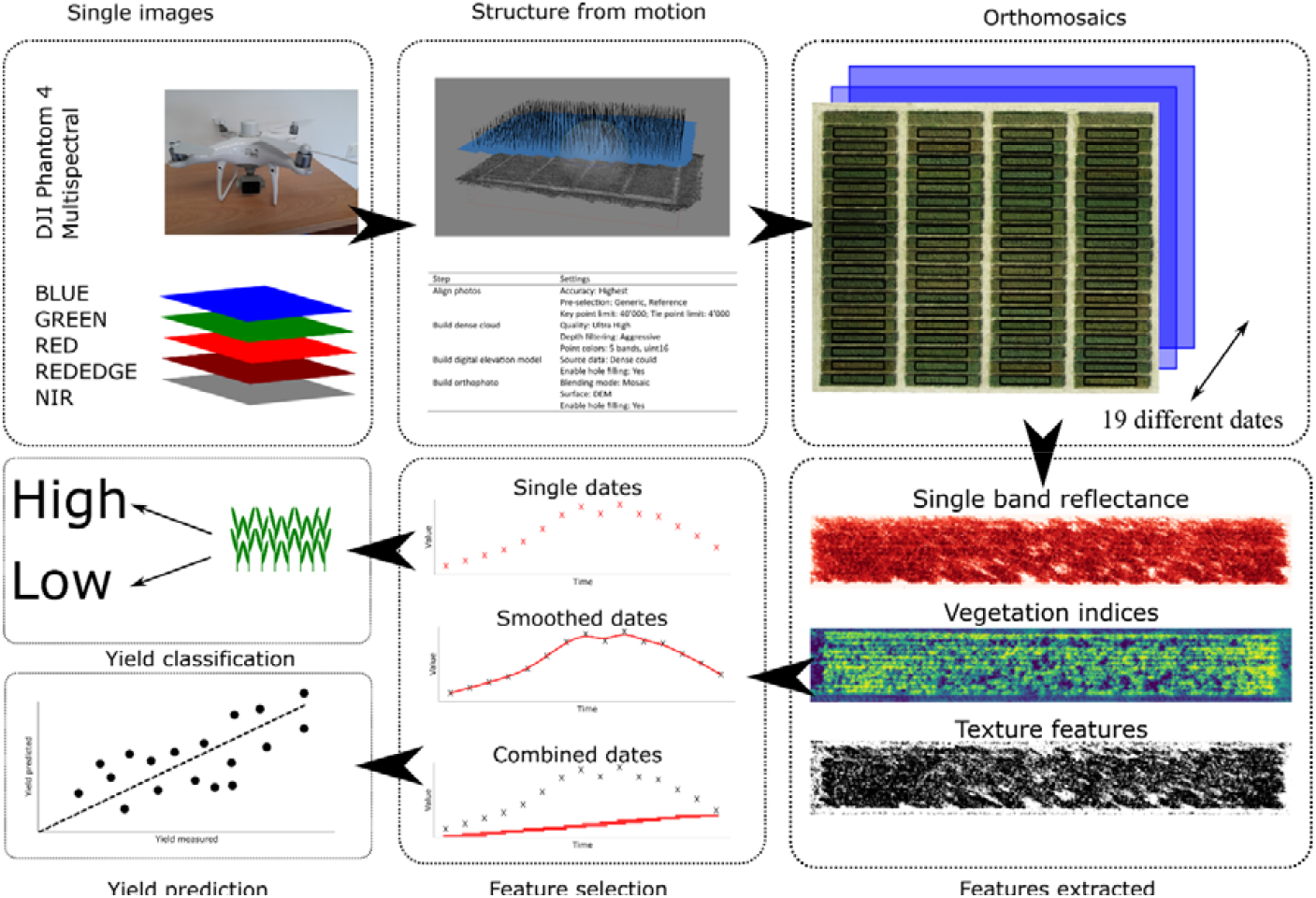
Workflow applied.

#### 2.3.1 Selection and calculation of spectral indices

To compare our approach across a range of vegetation indices (VIs), we classified them into five main groups based on their calculation method and selected a representative VI for each group. The five groups were differential-type, simple-ratio type, normalized differential type, three-band type, and combination of two spectral indices type (Table 1). We calculated the indices using a custom Python 3.7 script (Python Software Foundation, https://www.python.org/) and computed the median value for each index over the regions of interest (ROIs) corresponding to the plots.

**Table 1:** Vegetation indices (VIs) calculated.

| Index type | Index | Formula | Reference |
| --- | --- | --- | --- |
| **Difference** | **DVI** | $840 - 668$ | (Shibayama et al., 1999) |
| **Ratio** | **RVI** | $\frac{840}{668}$ | (Shibayama et al., 1999) |
| **Normalized** | **NDRE** | $\frac{840 - 717}{840 + 717}$ | (Barnes et al., 2000) |
| **Three Band** | **MCARI** | $((717 - 668) - 0.2 * (717 - 560)) * (\frac{717}{668})$ | (Daughtry et al., 2000) |
| **Combination of indices** | **CCII** | $\frac{TCARI}{OSAVI}$ | (Haboudane et al., 2002) |
| | **TCARI** | $3 * [(717 - 668) - 0.2 * (717 - 550) * (\frac{717}{670})]$ | |
| | **OSAVI** | $(1 + 1.16) * (\frac{840 - 668}{840 + 717 + 0.16})$ | |

#### 2.3.2 Selection and calculation of texture features

To generate a manageable number of TFs, we focused on calculating TFs for the RED reflectance band only as it had the best performance for yield prediction when using single bands. A 5 × 5 kernel size was used to calculate the GLCM features over the entire raster. This small kernel size was chosen because wheat leaf sizes are relatively small compared to our GSD. A quantization level of 32 was used, with the lowest level corresponding to the first percentile of the respective raster and the highest level corresponding to the 99th percentile. This ensured that we could still capture the variation in our image. GLCMs were constructed with a moving distance of 1 pixel and moving directions of 0°, 45° and 90° to eliminate possible effects of direction. The Contrast, Correlation, Dissimilarity, Energy, and Homogeneity features were extracted from each GLCM (Haralick et al., 1973) and saved as the center pixel in a raster. From these rasters, the final value per plot was extracted by averaging all values within the ROI. All calculations were performed using a custom Python 3.7 script (Python Software Foundation, https://www.python.org/). The extracted features are listed in (Table 2).

**Table 2:**
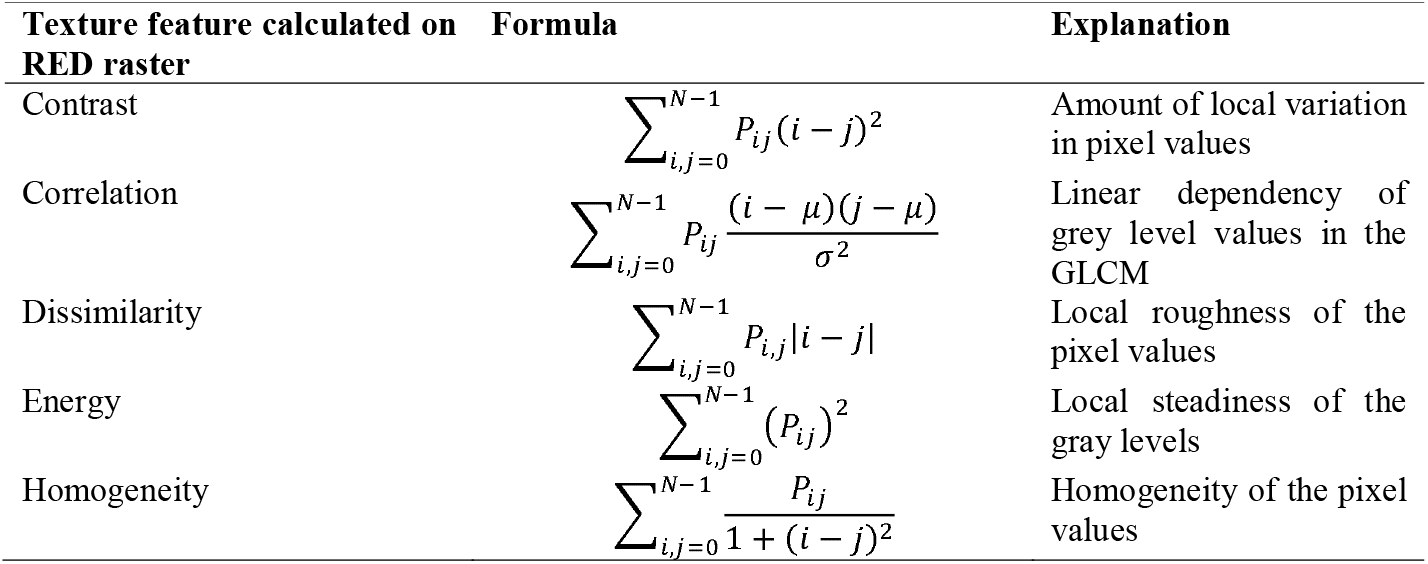
Calculation of grey correlation matrix features according to Haralick et al. (1973).

#### 2.3.3 Temporal processing of the extracted features

Three temporal feature selection strategies were evaluated (Figure 1). The first strategy involved selecting data from individual dates, resulting in one feature per observation. The second strategy involved smoothing the values per plot using splines, implemented in the statgenHTP package, with the default settings (Millet et al., 2022). Summed GDD from harvest were used as the time axis. Finally, features were selected using a moving time window with a width of 3. For each recorded date, the model included features from the current date and the previous as well as the following date, resulting in a total of three features per observation. This strategy is referred to as the moving window model.

### 2.4 Yield prediction model and yield potential classification model

To predict yield on a plot level and classify yield performance groups, we employed Random Forest (RF) machine learning models in R 4.2 (R Core Team, 2021). We optimized the number of trees per forest to 500 and used the R package *caret* (Kuhn, 2008). The number of trees per forest was set to 500 and the number of features per node was optimized by minimizing the root mean square error for the regression models and the accuracy for the classification models.

### 2.5 Statistical analysis

Pearson correlation coefficient between yield and spectral features was calculated using measurements taken on the 25.06.2021. At this date, most varieties were in the mid to end flowering and the correlation of VIs and yield was maximal for most VIs. The performances of the regression RF models were assessed by the coefficient of determination (R^2^) as well as the root mean square error (RMSE) using a 10-fold cross validation that was repeated 3 times and averaged:

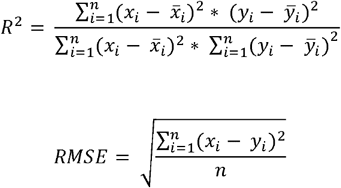

Where *x*_*i*_ and *y*_*i*_ represent the observed and the predicted yield, 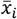 and 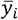 represent the mean of the observed and the predicted yield, respectively. *n* represents the number of samples. The performances of the classification RF models were assessed by the accuracy of the prediction using a 10-fold cross validation that was repeated 3 times and averaged:

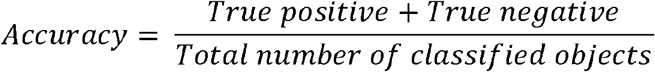

## 3 Results

### 3.1 Yield LAI and phenology

Substantial grain yield (GY) variation was observed between experimental plots (Table 3). The highest yield was observed in the variety *RGT-Reform* (411.4 g m^-2^), the highest yield in the variety *Skyfall* (642.7 g m^-2^). The high yielding varieties showed a significantly higher LAI during the stem elongation, the booting and at the late grain filling stage than the low yielding varieties (Figure 2). Phenology showed only few significant differences between the high and low yield groups, namely at the stem elongation and the flowering stage (p < 0.05). Still, it can be observed that the high yielding varieties were generally advanced in their phenology (Figure 2).

**Table 3:** Grain yield and phenology of the single varieties. Values represent the mean of the four replicates; the values in brackets represent the standard deviation of the four replicates.

| Variety | Grain Yield | Stem Elongation | Booting | Heading | Flowering | Early Grain Filling | Late Grain Filling |
| --- | --- | --- | --- | --- | --- | --- | --- |
| Absalon | 532,0 (32,8) | 635,0 (27,7) | 917,5 (37,3) | 1046,5 (23,6) | 1153,3 (38,7) | 1397,0 (59,2) | 1653,3 (79,2) |
| Aurelius | 541,9 (56,4) | 609,3 (3,5) | 963,5 (52,6) | 1069,3 (17,6) | 1109,0 (0,0) | 1368,8 (62,5) | 1673,3 (63,9) |
| Axioma | 473,0 (41,8) | 607,5 (4,0) | 950,3 (31,7) | 1042,8 (23,8) | 1121,5 (11,2) | 1350,0 (51,8) | 1636,5 (57,7) |
| Bernstein | 522,3 (33,7) | 626,8 (16,6) | 981,5 (6,9) | 1091,0 (30,2) | 1201,0 (70,6) | 1434,8 (27,3) | 1684,8 (93,6) |
| Bologna | 490,7 (33,3) | 625,5 (16,7) | 926,8 (61,1) | 1038,5 (30,0) | 1109,0 (0,0) | 1363,0 (69,7) | 1651,5 (64,4) |
| CH-Nara | 559,4 (30,2) | 635,0 (27,7) | 921,3 (16,7) | 1042,0 (16,2) | 1137,0 (30,9) | 1356,3 (47,8) | 1601,5 (23,7) |
| Chevignon | 598,5 (63,5) | 615,5 (30,6) | 911,8 (17,7) | 1050,8 (25,4) | 1144,8 (21,1) | 1358,3 (45,9) | 1579,3 (13,0) |
| Costello | 478,6 (38,8) | 635,5 (22,3) | 959,0 (31,2) | 1103,0 (38,0) | 1207,8 (67,8) | 1411,8 (39,7) | 1739,8 (2,5) |
| Dagmar | 617,1 (45,4) | 611,0 (0,0) | 969,0 (41,6) | 1052,0 (22,3) | 1114,0 (16,0) | 1345,3 (53,3) | 1653,5 (74,5) |
| Elixer | 540,4 (76,2) | 643,0 (27,7) | 953,0 (43,2) | 1118,3 (34,7) | 1219,5 (87,4) | 1444,0 (20,8) | 1742,0 (5,8) |
| Hyvento | 576,2 (51,1) | 641,3 (21,0) | 956,0 (34,8) | 1121,8 (6,0) | 1181,8 (41,7) | 1418,5 (23,7) | 1756,0 (119,6) |
| Julie | 544,2 (95,9) | 635,0 (27,7) | 915,3 (21,8) | 1057,3 (15,8) | 1142,8 (16,8) | 1347,3 (44,6) | 1648,5 (65,7) |
| Julius | 443,3 (14,6) | 639,0 (17,3) | 962,0 (26,9) | 1092,0 (40,1) | 1201,3 (65,4) | 1442,8 (28,5) | 1782,0 (62,2) |
| Montalbano | 584,1 (68,8) | 612,8 (3,5) | 954,0 (29,5) | 1104,0 (14,6) | 1192,0 (24,0) | 1428,0 (18,9) | 1720,5 (78,5) |
| Mv Nador | 552,4 (45,0) | 643,0 (27,7) | 962,3 (62,6) | 1084,3 (40,4) | 1157,3 (76,5) | 1343,8 (47,7) | 1624,5 (74,4) |
| Nogal | 504,2 (81,6) | 633,3 (29,9) | 972,8 (51,7) | 1058,8 (27,3) | 1084,0 (7,7) | 1281,5 (26,4) | 1602,0 (16,9) |
| RGT-Reform | 411,4 (60,2) | 629,0 (20,8) | 941,0 (14,3) | 1105,0 (9,9) | 1179,0 (48,3) | 1426,8 (18,9) | 1641,3 (69,7) |
| Rumor | 543,9 (15,5) | 623,0 (24,0) | 926,8 (22,5) | 1045,3 (14,5) | 1124,8 (23,2) | 1403,8 (10,5) | 1635,5 (67,6) |
| Skyfall | 642,7 (49,6) | 658,7 (82,6) | 909,0 (41,2) | 1067,8 (34,2) | 1157,5 (32,8) | 1424,8 (21,0) | 1642,8 (84,7) |
| All | 535,5 (73,8) | 628,5 (26,4) | 945,1 (39,3) | 1073,2 (35,8) | 1154,6 (54,3) | 1386,6 (56,6) | 1666,8 (80,1) |

**Figure 2:**
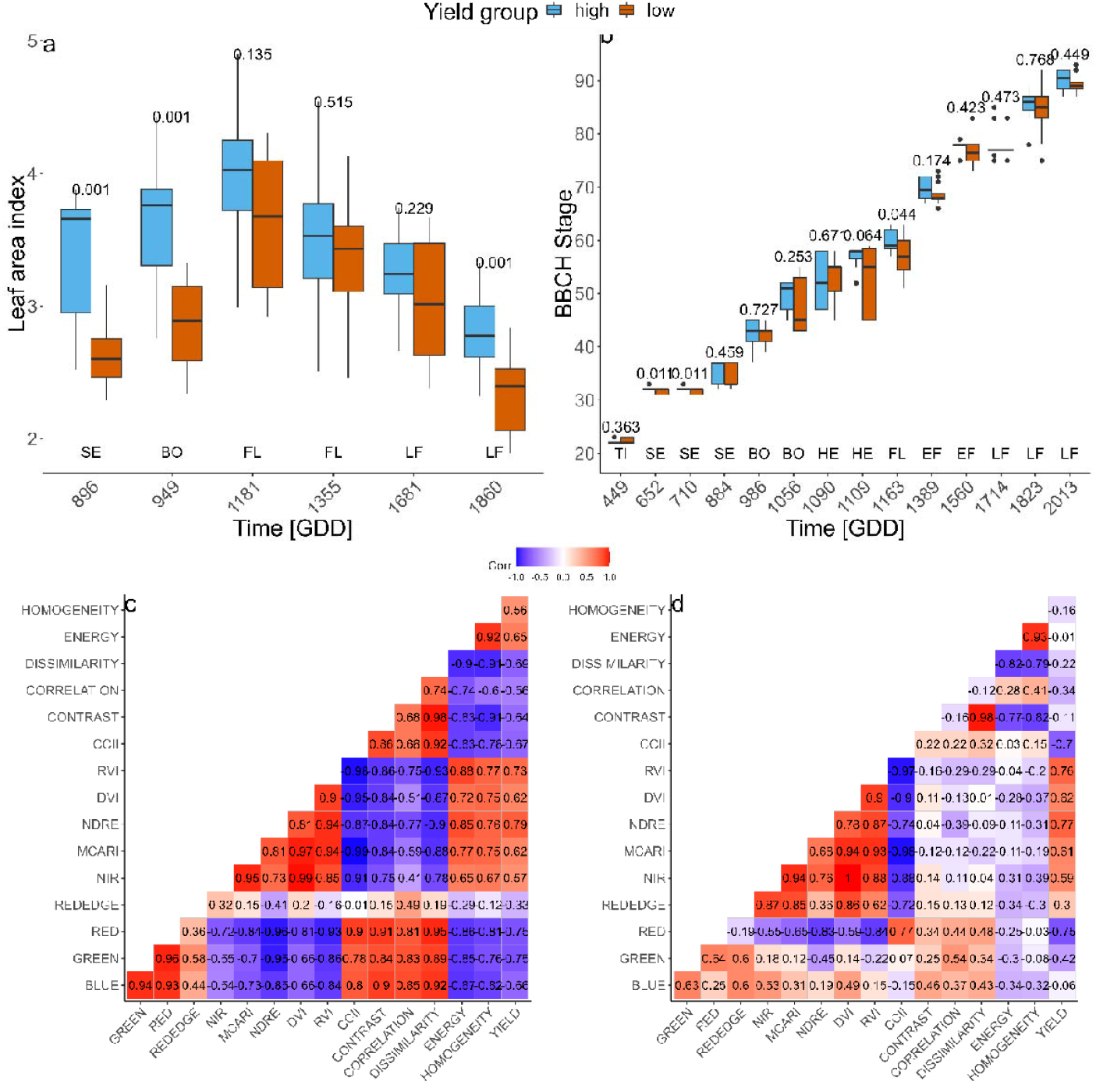
Leaf area index (a) and phenology (b) at different growth stages for the two high and low yield groups. Numbers above the boxplot pairs show the p-value of a t-test. Correlation of yield and indices on a single date at **949 GDD** after sowing at the booting stage (c) and **1389 GDD** after sowing at the beginning of the early grain filling stage (d). The numbers display the Pearson correlation coefficients.

### 3.2 Correlations between grain yield, the UAV based reflectance, vegetative indices and texture features

During booting, most extracted features show a high correlation with each other as well as with yield (Figure 2). The GREEN and RED reflectance showed an equally high correlation with yield (r = -0.75) at the booting stage, which was similar for RED at the early grain filling stage but significantly lower for the GREEN reflectance (r = -0.42). Among single band reflectance, the REDEDGE region expressed the lowest correlation with yield (r = -0.33) and was generally low correlated to other features.

Among VIs, the NDRE showed the highest correlation with yield during the booting stage (r = 0.79), followed by the RVI (r = 0.73), the CCII was negatively correlated to yield (r = -0.67) The correlations of VIs to yield do not change significantly at the early grain filling stage. TFs showed a moderate correlation with yield, and the DISSIMILARITY and ENERGY were found to be best two, respectively, with r values of -0.69 and 0.65 at the booting stage. Correlation of the TFs to yield changed drastically at the early grain filling stage, when the highest correlated feature CORRELATION showed an R of -0.34. The feature ENERGY was not correlated to yield at this stage anymore. Generally, VIs were the feature type that showed the highest correlation to yield at the booting as well as at the early grain filling stage.

### 3.3 Time series of UAV based reflectance, vegetative indices, texture features

Reflectance of the BLUE, GREEN and the RED band decreased with plant growth during tillering and stem elongation stages and increase with senescence, yielding a minimal reflectance around booting and flowering stages (Figure 3). The reflectance of the REDEDGE band was characterized by an increase in the first four measurement dates during tillering, a decrease during stem elongation, an increase at flowering and again a decrease with early grain filling stage. The reflectance of the NIR band increases with time to a maximum at late flowering stage and decreases again until full maturity. Significant differences were found for all reflectance bands, except REDEDGE, from the tillering to the early grain filling stage.

**Figure 3:**
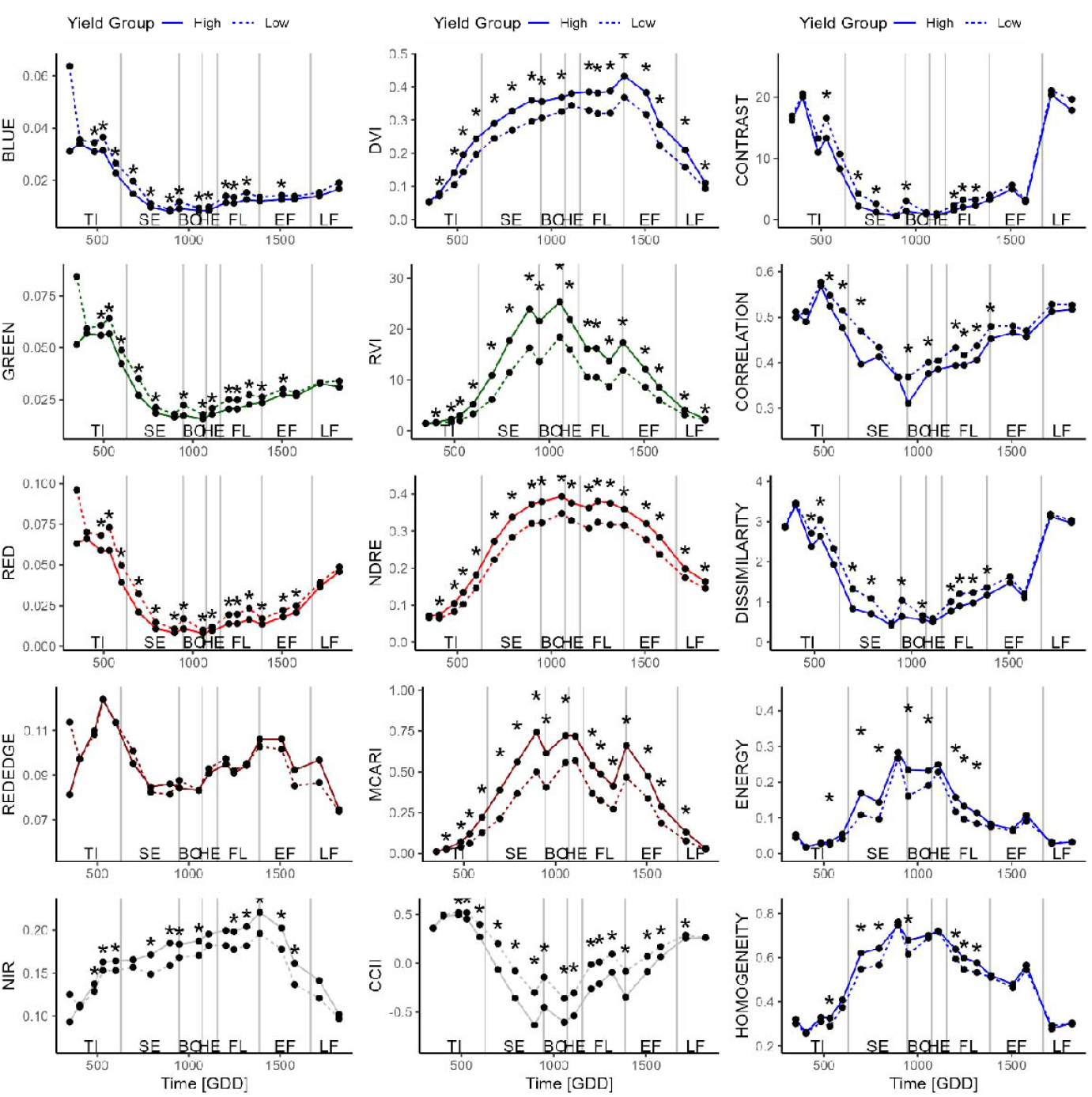
Dynamics of single band reflectances (left), vegetation indices (middle) and texture features (right) for different dates. The solid line shows the high yield group, the dashed line the mean value for the low yield group. The asterisks display significant differences after a t-test (p < 0.05) in the respective values and dates between the two yield groups.

All VIs showed significant differences between yield groups for all dates, except the first two and the last dates. DVI, RVI and NDRE even show significant differences between groups for all dates, except the first one (Figure 3).

The TFs CONTRAST, CORRELATION and DISSIMILARITY decreased during the stem elongation stage, reached a minimum around flowering stage and increase afterwards (Figure 3). ENERGY and

HOMOGENEITY showed an increase until heading stage and a steady decreased from then. Significant differences in the TFs between yield groups were mainly found from the tillering to the flowering stages but not later (Figure 3).

### 3.4 RF regression model for yield prediction using individual flights and time series of UAV traits

Significant effects on yield prediction model performance were found among the features chosen as well as the time points selected (Figure 4, Table 1).

**Figure 4:**
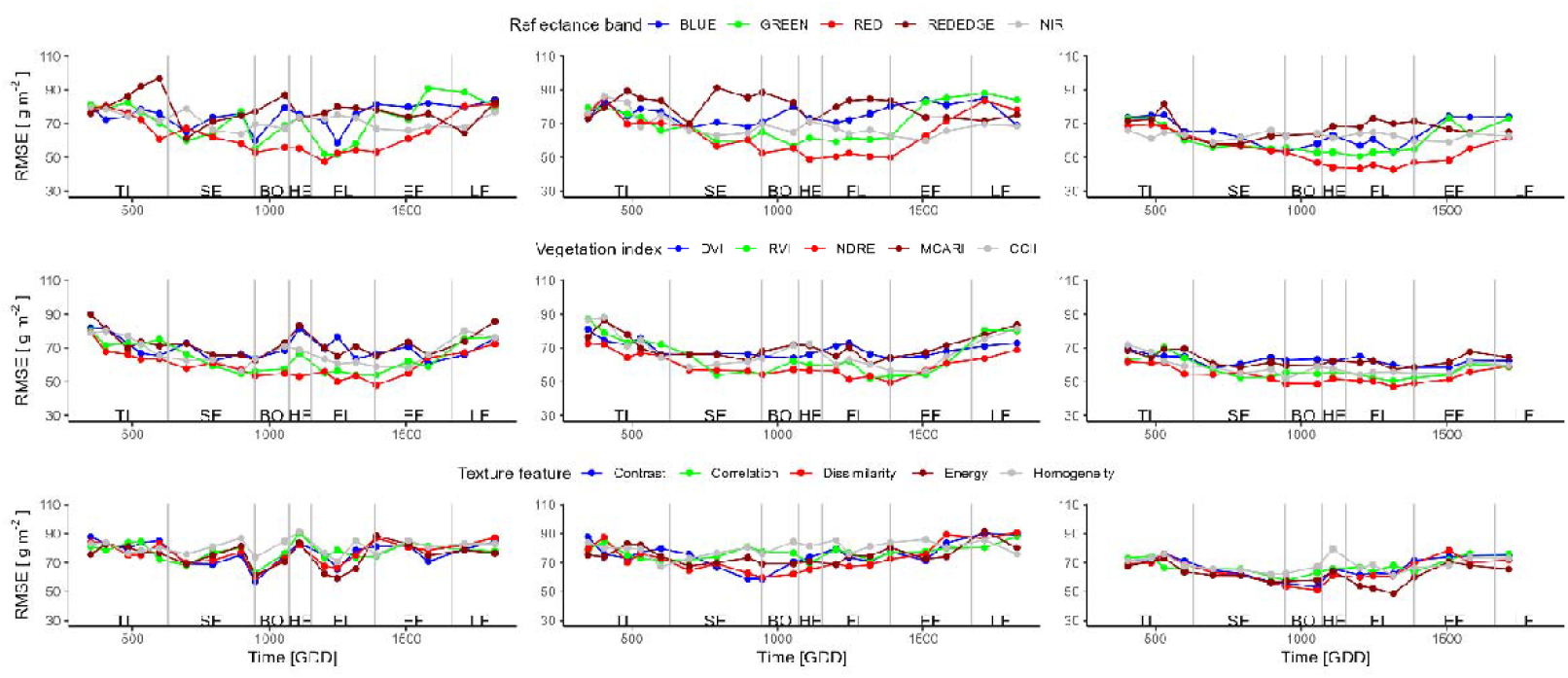
Root mean square errors of yield prediction modes built using single band reflectionsreflectances (top), vegetation indices (middle) and texture features (bottom). The models were built using measurements from single dates (left), smoothing features on single plots (middle) and choosing a window of three adjacent time points (right).

Based on reflectance measurements on a single date, the RED band showed the best performance in yield, with an RMSE of 47.4 g m^-2^ when taking the measurement at flowering stage, followed by the NDRE predicting yield with a RMSE of 47.9 g m^-2^ at the early grain filling stage. The performance of the prediction fluctuates significantly for single dates especially for single bands such as GREEN and BLUE, much less fluctuation can be observed for VIs. The best TF on a single day is CONTRAST predicting yield with an RMSE of 57.2 g m^-2^ at the booting stage. The performances of prediction models using TFs single dates are changing not only with phenology but also show big differences between two adjoining flights. For all TFs are affected in a similar way of single flights with eg. the measurement between the booting and the heading stage showed a better performance in yield prediction compared to the two adjacent dates (Figure 4).

Predicting yield from single dates that were previously smoothed on a plot level, was generally worse than predictions from non-smoothed single dates (Table 4). The RED band predicted yield with an RMSE of 49.0 g m^-2^ at heading stage. The best performing date for a single band often changed substantially after smoothing. Optimal time point for yield prediction using the NIR band was at stem elongation if the data is not smoothed and at early grain filling if the data is smoothed. Prediction of yield using the NIR band is slightly more accurate when using the smoothed data (RMSE of 59.9 g m^-2^) compared to using the original data (RMSE of 63.9 g m^-2^). The performance or optimal time points of VIs for yield prediction do not change significantly when smoothing the data. Smoothing of TFs improves or worsens the prediction of yield. It improves the prediction slightly for HOMOGENEITY but also worsens the prediction for ENERGY from an RMSE of 59.3 g m^-2^ to 67.7 g m^-2^. The fluctuations between dates became much less compared to non-smoothed single dates, especially in the highly fluctuating TFs.

**Table 4:** Display of the time points yielding the lowest RMSE for yield prediction for all reflectance bands, vegetation indices and texture features. Values in bold highlight the best model for a given spectral and temporal feature type combination.

| Raster | R<sup>2</sup> | RMSE | Date | PS | VI | R<sup>2</sup> | RMSE | Date | PS | TF | R<sup>2</sup> | RMSE | Date | PS |
| --- | --- | --- | --- | --- | --- | --- | --- | --- | --- | --- | --- | --- | --- | --- |
| Single time points |  |  |  |  |  |  |  |  |  |  |  |  |  |  |
| BLUE | 0.45 | 58.8 | 1248 | FL | DVI | 0.38 | 60.8 | 1580 | EF | **CONTRAST** | **0.46** | **57.2** | **949** | **BO** |
| GREEN | 0.57 | 51.8 | 1248 | FL | RVI | 0.52 | 54.1 | 1389 | EF | CORRELATION | 0.37 | 63.1 | 949 | BO |
| **RED** | **0.63** | **47.4** | **1202** | **FL** | **NDRE** | **0.63** | **47.9** | **1389** | **EF** | DISSIMILARITY | 0.42 | 60.0 | 949 | BO |
| REDEDGE | 0.39 | 61.3 | 697 | SE | MCARI | 0.38 | 62.6 | 949 | BO | ENERGY | 0.43 | 59.3 | 1248 | FL |
| NIR | 0.36 | 63.9 | 896 | SE | CCII | 0.48 | 56.1 | 896 | SE | HOMOGENEITY | 0.28 | 70.4 | 1248 | FL |
| Smoothed single time points |  |  |  |  |  |  |  |  |  |  |  |  |  |  |
| BLUE | 0.37 | 67.4 | 696 | SE | DVI | 0.30 | 63.7 | 1388 | EF | CONTRAST | 0.41 | 58.7 | 949 | BO |
| GREEN | 0.48 | 56.6 | 1055 | BO | RVI | 0.50 | 52.3 | 1317 | FL | CORRELATION | 0.21 | 69.8 | 1110 | HE |
| **RED** | **0.58** | **49.0** | **1110** | **HE** | **NDRE** | **0.57** | **49.6** | **1388** | **EF** | **DISSIMILARITY** | **0.42** | **59.9** | **949** | **BO** |
| REDEDGE | 0.32 | 69.9 | 696 | SE | MCARI | 0.37 | 60.2 | 1317 | FL | ENERGY | 0.26 | 67.7 | 696 | SE |
| NIR | 0.41 | 59.9 | 1509 | EF | CCII | 0.45 | 56.0 | 1509 | EF | HOMOGENEITY | 0.29 | 67.2 | 600 | TI |
| Moving time window |  |  |  |  |  |  |  |  |  |  |  |  |  |  |
| BLUE | 0.54 | 53.3 | 1316 | FL | DVI | 0.43 | 58.1 | 697 | SE | CONTRAST | 0.51 | 53.2 | 1056 | BO |
| GREEN | 0.58 | 50.7 | 1202 | FL | RVI | 0.56 | 50.6 | 1316 | FL | CORRELATION | 0.41 | 57.6 | 949 | BO |
| **RED** | **0.69** | **42.7** | **1316** | **FL** | **NDRE** | **0.64** | **47.0** | **1316** | **FL** | DISSIMILARITY | 0.54 | 51.1 | 1056 | BO |
| REDEDGE | 0.42 | 57.8 | 697 | SE | MCARI | 0.44 | 57.2 | 1316 | FL | **ENERGY** | **0.60** | **48.7** | **1316** | **FL** |
| NIR | 0.42 | 59.1 | 697 | SE | CCII | 0.56 | 51.7 | 949 | BO | HOMOGENEITY | 0.36 | 61.1 | 1316 | FL |

Based on reflectance measurements across several dates, using each of the individual bands improved the yield prediction. The RED band, which already showed the best performance for single date yields an RMSE of 42.7 g m^-2^ when selecting three dates around flowering. The prediction of yield using VIs also improved for all bands but to less extent as for the single reflectance bands. The prediction using NDRE improved slightly to an RMSE of 47.0 g m^-2^. TFs were the feature class that improved most when using several time points for yield prediction and all features were better able to prediction yield compared to single dates. A substantial improvement was found in the ENERGY feature that was able to predict yield with an RMSE of 48.7 g m^-2^ when using measurements around flowering stage (Table 4). Using a time window lead to predictions that were much less fluctuating compared to measurements on single flights (Figure 4). Predictions using the RED band show am underestimation of yields higher than 630 g m^-2^ (Figure 6).

### 3.5 RF classification model for classifying the low and high yielding varieties using individual flights and time series of UAV traits

Classification of yield groups using single bands was best with the RED band resulting in an accuracy of 0.962 followed by the BLUE band with an accuracy of 0.873, both at the flowering stage. The NIR band performed best during the booting stage. Classification using VIs was lower than accuracy for single bands and the highest accuracy was found for the RVI VI (0.875). All VIs show their best performance at the heading stage or later. The TFs showed a similar performance for yield type classification as the VIs. The best performing TF was CONTRAST with an accuracy of 0.898 when being measured during stem elongation stage. Fluctuations in accuracy between subsequent flights was relatively high for models built using single band reflectance as well as models using TFs on single dates (Figure 5).

**Figure 5:**
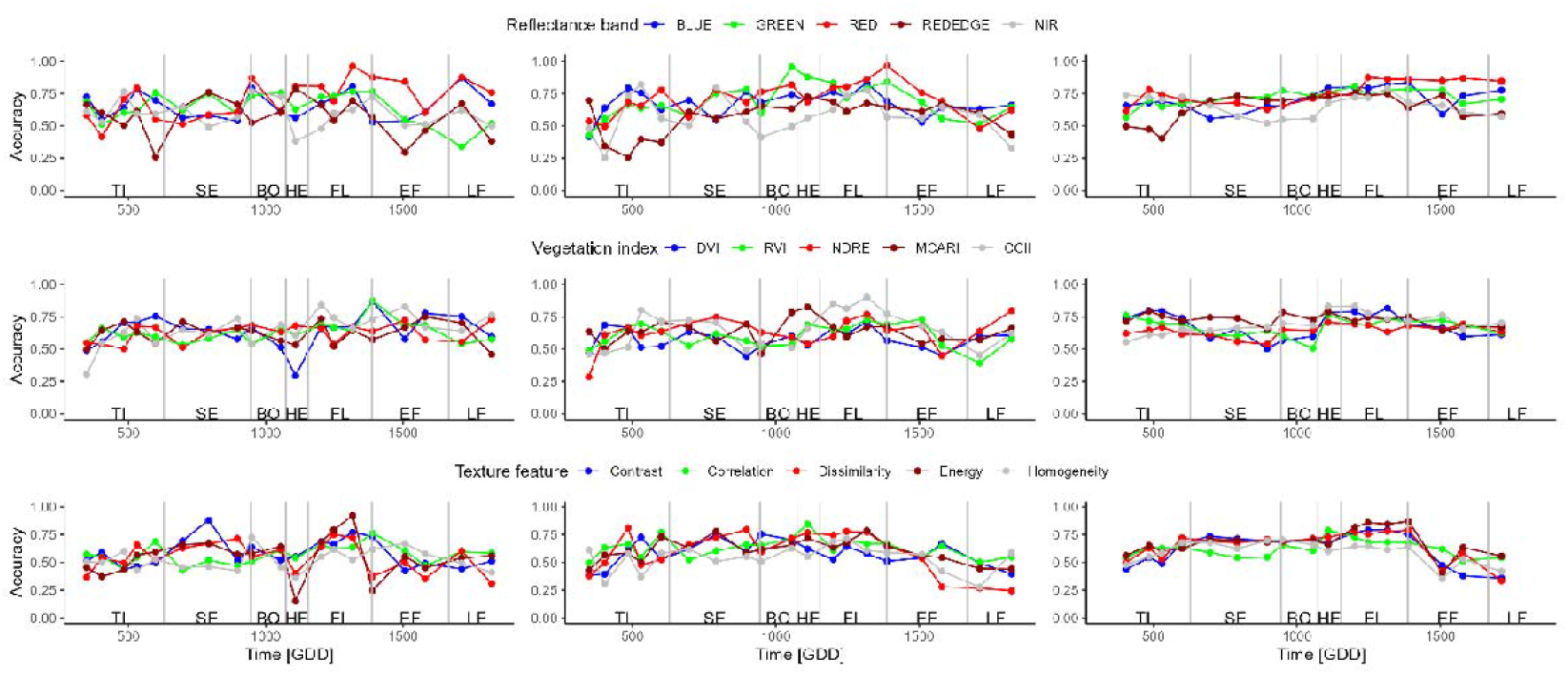
Accuracy prediction models built using single band reflectances (top), vegetation indices (middle) and texture features (bottom). The models were built using measurements from single dates (left), cumulating features from the beginning (middle) and choosing a window of three time points (right).

**Figure 6:**
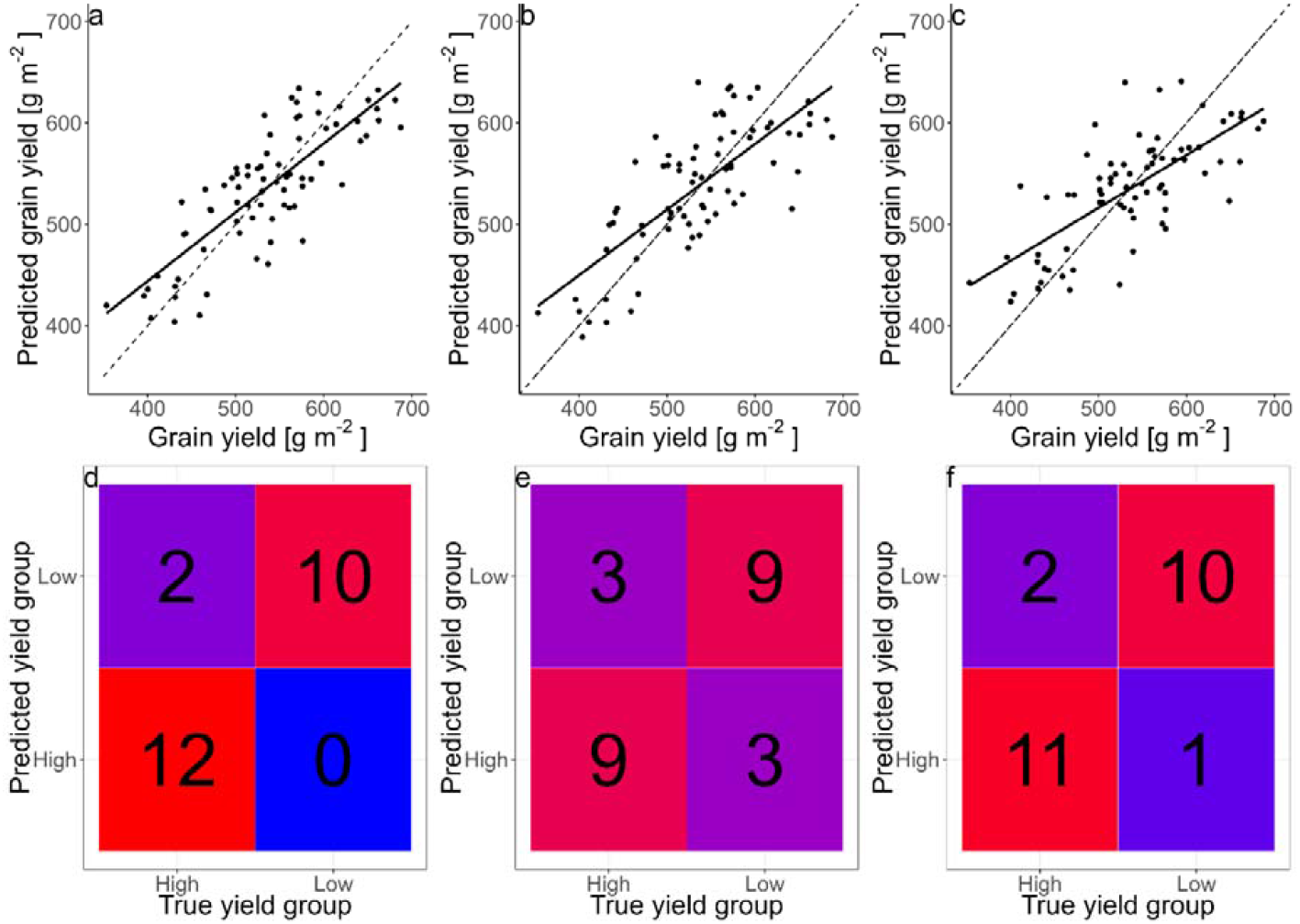
Yield prediction based on the red band (a), the NDRE index (b) and the energy TF (c) from moving window features at the best performing dates (Table 4). Confusion matrices of yield type predictions based on data of the best performing single dates of the RED band (d), the CCII VI (e) and the energy TF (f).

Smoothing reflectances did not change improve the accuracy for yield type classification for the RED band, yielding an accuracy of 0.963 on time point later at the early grain filling stage (Table 5). The accuracy of the GREEN improved substantially to 0.955 at the booting stage and the NIR band increase in accuracy as well, while a date on the tillering stage was found to be most suitable. Accuracies of the DVI and RVI VIs dropped whereas the accuracies increase slightly for the other VIs after smoothing. Eg. the accuracy of the CCII increased to 0.902 and was achieved during flowering stage. The best performing TF ENERGY decreased when smoothing to 0.787. CORRELATION however was better able to classify yield types with an accuracy of 0.845. DISSIMILARITY showed the best performance at the tillering stage after smoothing. Smoothing changed the fluctuations in accuracy between single flights slightly for the VIs and to a bigger extent for the single band reflectances and mainly for the TFs (Figure 5).

**Table 5:**
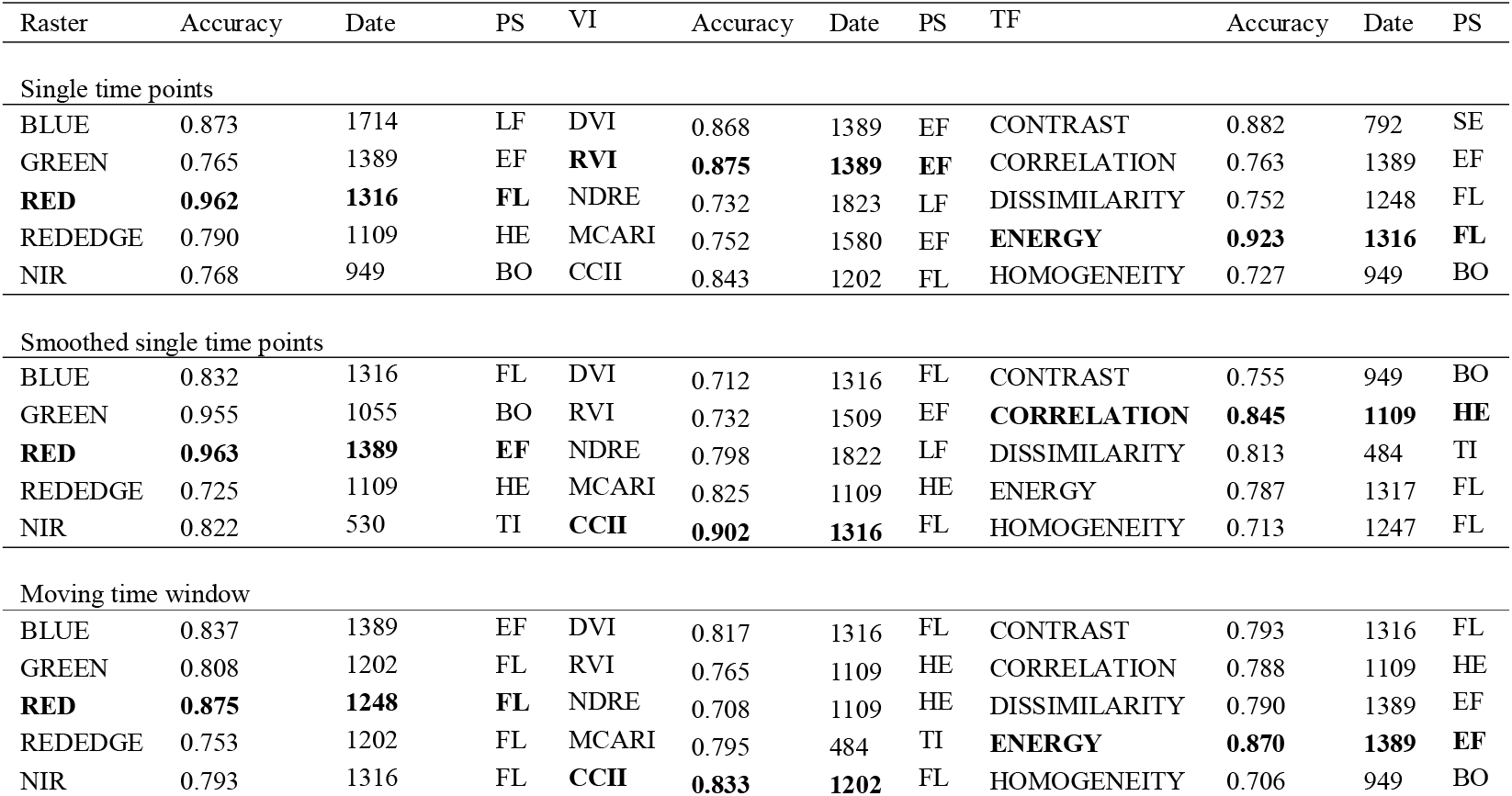
Display of the classification accuracy of different yield groups using predictors for all reflectance bands, vegetation indices and texture features at all dates.

The moving time window did not improve yield type classification but worsened for the RED band yielding an accuracy of 0.875 and did not change the accuracies of the other single bands significantly (Table 5). A similar result can be found for the moving window when applied to the VIs where the RVI yielded an accuracy of 0.765 at the flowering stage. A moving time window improved the classification for all VIs except DVI. However, these changes were only small for all VIs. The best VI for yield prediction using a moving window is the RVI with an accuracy of 0.856. Predictions by TFs worsen when using several time points as features compared to single dates. Using a time window, the

TF ENERGY and DISSIMILARITY show almost similar accuracies of 0.803 and 0.801, respectively. While the accuracies were not significantly improved by using a time window for prediction, the fluctuations between dates were reduced significantly and thus the performances of the predictions became much more stable among dates (Figure 5).

## 4 Discussion

### 4.1 Dynamic responses of individual bands

The red edge bands have been widely studied for assessing crop performance and yield in various crops, including wheat (Horler et al., 1983). Canopy reflectance in the red edge wavelength range is influenced by two optical properties of canopies: chlorophyll absorption in the red region and multiple scattering effects on the near-infrared (NIR). Although red edge reflectance has been commonly used for yield prediction in previous studies, our findings suggest that this may not be a reliable indicator of yield in our specific case.

The inter-variety variability in canopy reflectance in the red-edge region might be large and have overweight the subtle differences in canopy reflectance associated with yield, making the red edge reflectance less useful for yield prediction. In addition, the correlation of reflectance in the rededge region with yield is known to change quickly with the exact wavelength measured in the red edge region (Pavuluri et al., 2015). In contrast, visible bands (Blue, Green and Red) can be more sensitive to yield-related variations in chlorophyll, biomass accumulation during the tillering until the and the stem elongation stage until the beginning of the booting stage since they are known to be correlated to both, chlorophyll concentration and LAI (Daughtry et al., 2000). Accordingly, our results showed that the RED, GREEN, and BLUE bands were among the most effective spectral features for yield prediction, exhibiting significant differences between high and low yielding varieties at almost all measurement dates. These visible bands are highly sensitive to chlorophyll, with their reflectance decreasing during the transition from the stem elongation to the beginning of the booting stage, and increasing again from heading until harvest due to senescence and chlorophyll degradation. Therefore, our findings suggest that the flowering stage in which the chlorophyll content reaches the peak is crucial.

The NIR region is known to be sensitive to leaf area and especially ground cover (Korobov & Railyan, 1993), making it a useful band for predicting biomass and yield. Our results indicated that the NIR band performed best during the stem elongation stage for yield prediction and at the booting stage when there were significant differences in leaf area index (LAI) between the two yield groups. This aligns with the findings by Korobov and Railyan (1993), who reported a higher correlation of NIR reflectance with dry matter and ground cover during booting stage compared to later stages. Thus, normalizing the difference of the NIR and the REDEDGE reflectance in form of the NDRE index, showed a good performance for chlorophyll estimation (Barnes et al., 2000). Usually, VIs containing information from the red edge region of the spectrum are considered being more sensitive to chlorophyll absorption in dense canopies (Nguy-Robertson et al., 2012). It is expected that combining the highly LAI-sensitive NIR band with the red edge band containing more information about leaf pigments in the canopy and therefore improves the performance of our yield prediction model at the flowering to early grain filling stages.

### 4.2 The influence of growth stage for yield prediction and classification

The performance of yield prediction and classification depends highly on the growth stages of the crop. Our study found that the flowering stage and early grain filling stage are the most suitable for predicting yield in winter wheat. This is consistent with the findings of other studies (Hassan et al., 2019; Prey et al., 2022; Prey et al., 2020; Wang et al., 2022). It is often argued that yield cannot be measured directly using remote sensing approaches, given the fact that yield formation in wheat involves several components, including ear density, kernel number per ear, and grain weight (Satorre & Slafer, 1999). Furthermore, these components form at different stages and are therefore not present at every phenological stage. Still, remotely sensed information and variations in canopy images and reflectance are often closely related to these yield components, such as vegetative biomass and chlorophyll content (Wu et al., 2008). Therefore, when these traits usually reach their peak levels around the flowering stage, i.e., the crop transitions from vegetative to generative growth, their associated variations in canopy reflectance also indicate the variability in yield

Early differences in biomass and LAI dynamics between genotypes are well-documented (Grieder et al., 2015; Pang et al., 2014), and these differences can be useful for predicting yield and classifying varieties in early growth stages. While our study found that yield prediction at the tillering stage feasible, it remains to be challenging for excluding the poorly-performing varieties. From tillering to harvest, wheat is known to compensate for e.g. a low stand count by altering the number of yield components (Holen et al., 2001) complicating predictions at this stage. This might be the reason, that our classification models were more performant in detecting the high yielding varieties. It can be argued that our approach has a limitation that we cannot be certain whether the algorithm is sensitive to canopy traits that are yield-dependent or to variety-specific traits that do not influence yield performance. Therefore, the validation of our models in other varieties or environments will be critical. After flowering, the performance of our models then decreased with the onset of senescence. While yield formation does continue during senescence (Anderegg et al., 2020; Spano et al., 2003), our results showed that this stage is less correlated with grain yield than the earlier stages. Further, no significant differences in phenology were detected between the yield groups, suggesting that the yield differences between the high- and low yield groups are not likely due to their senescence dynamics. Unlike in our study employing an identical/moderate fertilization rate, even under high N conditions, differences in the onset of senescence were not found to be significantly correlated to grain yield per unit crop N uptake at harvest (Gaju et al., 2011) making classification at this time point difficult. Nevertheless, our study confirms that the tillering stage is an already promising time for variety classification, as evidenced by the strong performance of our classification models at this stage.

### 4.3 Comparison of variable- and feature types for yield prediction and classification

Our study found that single-band reflectance, such as the RED band, were as effective as or even more effective than vegetation indices (VIs) for predicting yield. The RED band is known to be related to leaf area index (LAI), although this relationship is often nonlinear (Hinzman et al., 1984) and therefore requires non-linear methods such as RF to perform well for yield prediction. Pavuluri et al. (2015) found a saturation of RED reflectance when predicting yield, which can also be found in our prediction models. In contrast, VIs typically show good linear correlations with grain yield, with NDVI being widely used for yield prediction (Hassan et al., 2019). Many VIs have been screened by Prey et al. (2020) and few have been showing a consistent performance over the years, which makes a general selection difficult. Further, VIs narrow down the information that is accessible and Vatter et al. (2022) found good performances for yield prediction when using 11 wavebands from a multispectral camera that were fed to a deep learning model. Machine learning approaches based on reflectance were found to not improve the accuracy yield prediction but showed great potential in predicting grain protein (Zhou et al., 2021). Machine learning models, however, often require a big amount of training data, which can be challenging to gather.

TFs are complex in their calculation and they offer a variety of possible ways of calculation, possible combination with underlying rasters and ways to be calculated. Detailed information on how TFs are calculated is often lacking (Wang et al., 2021; Zhang et al., 2021; Zheng et al., 2019). Therefore, TFs still have to be examined in detail and their parameters optimized under different experimental conditions and scenarios of sensing data collection. We calculated TFs in a standardized way, but still found a high variability between dates. TFs are further known to be highly dependent on the GSD and therefore the flight height (Zheng et al., 2019). Therefore, smoothing benefitted the yield prediction by making the performance more stable but not better. A novel approach is presented by Herrero-Huerta et al. (2020) who calculated so-called canopy roughness directly on the point cloud from the structure from motion processing and showed its correlation to biomass. TFs are further often used in combination because there might be additional information (Liu et al., 2022; Wang et al., 2021), especially in the later stage, when they are not strongly correlated to single band reflectances and VIs anymore as indicate by our results.

### 4.4 Limitations and outlook

The red-edge position and its shape is often used estimate the stress status of field crops (Boochs et al., 1990; Guyot et al., 1988). However, it is obvious that the dynamics (time series) of the Red-edge band is difficult to interpret compared to the visible bands. During the early stages of tillering, the red-edge reflectance increased, possibly due to an increase of ground cover, whereas later it decreased again, when the canopy height increased during the SE stage. At the beginning of heading another increase in the Red-edge reflectance can be observed, accompanied with the increase of reflectance in the visible bands. However, in contrast to other bands, the Red-edge reflectance decreases with the onset of senescence at the early grain filling stage, possibly due to a reduction in chlorophyll and shrinking canopy structure (Wang et al., 2022). However, fluctuation also occurs during the mentioned stable period. These fluctuations can be of various origins. For instance, the appearance of the canopy might change significantly due to the emergence of the spikes. Although this study was unable to exploit the entire shape of the red-edge reflectance, due to limitations in our multispectral camera having one band in the red edge region, future work should further advance the understanding of the dynamics of red-edge reflectance and responsible canopy characteristics.

Also, features should in addition to their performance for yield prediction be assessed regarding their heritability (h^2^) since breeders are interested in knowing the genetic variation underlying a trait or in our case a spectral feature. Generally, this study shows that a trait time series followed by smoothing and a moving window allow for more stable predictions when also not better predictions.

## 5 Conclusions

Most spectral and texture features derived from the canopy multispectral images were related to variations in yield, but they-delivered the best predictions of yield only between the booting and end of the flowering stage. However, in earlier stages, the visible (Red, Green, Blue) bands can accurately predict yields and distinguish between the low- and high-yielding genotypes. Single-band reflectance, particularly the red band, is a reliable predictor of yield. Combining additional bands, such as the red-edge and NIR, into VIs like NDRE, improves performance significantly, but it limits the machine learning algorithm’s ability to build a strong model. Texture features generally performed poorly for yield prediction, and their performances were inconsistent across dates in this study, suggesting that further research is still needed to better understand the applicability of different texture features for yield- and traits predictions and methods to optimize texture feature extraction. Smoothing or combining data across a time series can enhance the performance of yield prediction and classification models, particularly in the early growth stages. Future studies should combine different feature types to leverage complementary information captured by different types of multispectral features and variables.

## Author contributions

KY designed the experiment. MC managed the UAV flights for aerial imagery, analyzed the data, and conducted the ground-based field measurements with supervision from KY. MC and KY wrote and revised the manuscript. Both authors contributed to the article and approved the submitted version.

## Acknowledgements

We thank Juan Herrera (Agroscope, Switzerland) for composing the variety-testing panel and providing the seeds. Further, we thank Wolfgang Heer for trial management, Jürgen Plass for technical support with the spectral measurements and Michael Thaler for the help with data collection.

## References

Anderegg, J., Yu, K., Aasen, H., Walter, A., Liebisch, F., & Hund, A. (2020). Spectral Vegetation Indices to Track Senescence Dynamics in Diverse Wheat Germplasm [Original Research]. Frontiers in Plant Science, 10(1749). https://doi.org/10.3389/fpls.2019.01749

Araus, J. L., Buchaillot, M. L., & Kefauver, S. C. (2022). High Throughput Field Phenotyping. In M. P. Reynolds & H.-J. Braun (Eds.), Wheat Improvement: Food Security in a Changing Climate (pp. 495–512). Springer International Publishing. https://doi.org/10.1007/978-3-030-90673-3_27

Araus, J. L., & Cairns, J. E. (2014). Field high-throughput phenotyping: the new crop breeding frontier. Trends in Plant Science, 19(1), 52–61. https://doi.org/10.1016/j.tplants.2013.09.008

Barnes, E., Clarke, T., Richards, S., Colaizzi, P., Haberland, J., Kostrzewski, M., Waller, P., Choi, C., Riley, E., & Thompson, T. (2000). Coincident detection of crop water stress, nitrogen status and canopy density using ground based multispectral data. Proceedings of the Fifth International Conference on Precision Agriculture, Bloomington, MN, USA,

Boochs, F., Kupfer, G., Dockter, K., & KÜHbauch, W. (1990). Shape of the red edge as vitality indicator for plants. International Journal of Remote Sensing, 11(10), 1741–1753. https://doi.org/10.1080/01431169008955127

Bowman, B. C., Chen, J., Zhang, J., Wheeler, J., Wang, Y., Zhao, W., Nayak, S., Heslot, N., Bockelman, H., & Bonman, J. M. (2015). Evaluating Grain Yield in Spring Wheat with Canopy Spectral Reflectance. Crop Science, 55(5), 1881–1890. https://doi.org/https://doi.org/10.2135/cropsci2014.08.0533

Bukowiecki, J., Rose, T., Ehlers, R., & Kage, H. (2020). High-Throughput Prediction of Whole Season Green Area Index in Winter Wheat With an Airborne Multispectral Sensor [Original Research]. Frontiers in Plant Science, 10. https://doi.org/10.3389/fpls.2019.01798

Cabrera-Bosquet, L., Crossa, J., von Zitzewitz, J., Serret, M. D., & Luis Araus, J. (2012). Highthroughput Phenotyping and Genomic Selection: The Frontiers of Crop Breeding ConvergeF. Journal of Integrative Plant Biology, 54(5), 312–320. https://doi.org/https://doi.org/10.1111/j.1744-7909.2012.01116.x

Culbert, P. D., Pidgeon, A. M., St.-Louis, V., Bash, D., & Radeloff, V. C. (2009). The Impact of Phenological Variation on Texture Measures of Remotely Sensed Imagery. IEEE Journal of Selected Topics in Applied Earth Observations and Remote Sensing, 2(4), 299–309. https://doi.org/10.1109/JSTARS.2009.2021959

Daughtry, C. S. T., Walthall, C. L., Kim, M. S., de Colstoun, E. B., & McMurtrey, J. E. (2000). Estimating Corn Leaf Chlorophyll Concentration from Leaf and Canopy Reflectance. Remote Sensing of Environment, 74(2), 229–239. https://doi.org/https://doi.org/10.1016/S0034-4257(00)00113-9

Deutsche Landesvermessung. Satellitenpositionierungsdienst der Deutschen Landesvermessung. Retrieved 22.02.2023 from https://www.ldbv.bayern.de/produkte/gis_gps/dgps/dgps.html

Duan, T., Chapman, S. C., Guo, Y., & Zheng, B. (2017). Dynamic monitoring of NDVI in wheat agronomy and breeding trials using an unmanned aerial vehicle. Field Crops Research, 210, 71–80. https://doi.org/https://doi.org/10.1016/j.fcr.2017.05.025

Elsayed, S., Elhoweity, M., Ibrahim, H. H., Dewir, Y. H., Migdadi, H. M., & Schmidhalter, U. (2017). Thermal imaging and passive reflectance sensing to estimate the water status and grain yield of wheat under different irrigation regimes. Agricultural Water Management, 189, 98–110. https://doi.org/https://doi.org/10.1016/j.agwat.2017.05.001

Fernandez-Gallego, J. A., Kefauver, S. C., Vatter, T., Aparicio Gutiérrez, N., Nieto-Taladriz, M. T., & Araus, J. L. (2019). Low-cost assessment of grain yield in durum wheat using RGB images. European Journal of Agronomy, 105, 146–156. https://doi.org/https://doi.org/10.1016/j.eja.2019.02.007

Gaju, O., Allard, V., Martre, P., Snape, J. W., Heumez, E., LeGouis, J., Moreau, D., Bogard, M., Griffiths, S., Orford, S., Hubbart, S., & Foulkes, M. J. (2011). Identification of traits to improve the nitrogen-use efficiency of wheat genotypes. Field Crops Research, 123(2), 139–152. https://doi.org/https://doi.org/10.1016/j.fcr.2011.05.010

Grieder, C., Hund, A., & Walter, A. (2015). Image based phenotyping during winter: a powerful tool to assess wheat genetic variation in growth response to temperature. Functional Plant Biology, 42(4), 387–396. https://doi.org/https://doi.org/10.1071/FP14226

Guyot, G., Baret, F., & Major, D. (1988). High spectral resolution: determination of spectral shifts between the red and near infrared. ISPRS Congress,

Haboudane, D., Miller, J. R., Tremblay, N., Zarco-Tejada, P. J., & Dextraze, L. (2002). Integrated narrow-band vegetation indices for prediction of crop chlorophyll content for application to precision agriculture. Remote Sensing of Environment, 81(2), 416–426. https://doi.org/https://doi.org/10.1016/S0034-4257(02)00018-4

Haralick, R. M., Shanmugam, K., & Dinstein, I. (1973). Textural Features for Image Classification. IEEE Transactions on Systems, Man, and Cybernetics, SMC-3(6), 610–621. https://doi.org/10.1109/TSMC.1973.4309314

Hassan, M. A., Yang, M., Rasheed, A., Yang, G., Reynolds, M., Xia, X., Xiao, Y., & He, Z. (2019). A rapid monitoring of NDVI across the wheat growth cycle for grain yield prediction using a multi-spectral UAV platform. Plant Science, 282, 95–103. https://doi.org/https://doi.org/10.1016/j.plantsci.2018.10.022

Herrero-Huerta, M., Bucksch, A., Puttonen, E., & Rainey, K. M. (2020). Canopy Roughness: A New Phenotypic Trait to Estimate Aboveground Biomass from Unmanned Aerial System. Plant Phenomics, 2020, 6735967. https://doi.org/10.34133/2020/6735967

Hinzman, L., Bauer, M. E., & Daughtry, C. (1984). Growth and reflectance characteristics of winter wheat canopies.

Holen, D. L., Bruckner, P. L., Martin, J. M., Carlson, G. R., Wichman, D. M., & Berg, J. E. (2001). Response of Winter Wheat to Simulated Stand Reduction. Agronomy Journal, 93(2), 364–370. https://doi.org/https://doi.org/10.2134/agronj2001.932364x

Horler, D. N. H., Dockray, M., & Barber, J. (1983). The red edge of plant leaf reflectance. International Journal of Remote Sensing, 4(2), 273–288. https://doi.org/10.1080/01431168308948546

Hund, A., Kronenberg, L., Anderegg, J., Yu, K., & Walter, A. (2019). Non-invasive field phenotyping of cereal development. In Advances in breeding techniques for cereal crops (pp. 249–292). Burleigh Dodds Science Publishing.

Kirchgessner, N., Liebisch, F., Yu, K., Pfeifer, J., Friedli, M., Hund, A., & Walter, A. (2017). The ETH field phenotyping platform FIP: a cable-suspended multi-sensor system. Functional Plant Biology, 44(1), 154–168. https://doi.org/https://doi.org/10.1071/FP16165

Korobov, R. M., & Railyan, V. Y. (1993). Canonical correlation relationships among spectral and phytometric variables for twenty winter wheat fields. Remote Sensing of Environment, 43(1), 1–10. https://doi.org/https://doi.org/10.1016/0034-4257(93)90059-7

Kronenberg, L., Yates, S., Boer, M. P., Kirchgessner, N., Walter, A., & Hund, A. (2020). Temperature response of wheat affects final height and the timing of stem elongation under field conditions. Journal of Experimental Botany, 72(2), 700–717. https://doi.org/10.1093/jxb/eraa471

Kuhn, M. (2008). Building Predictive Models in R Using the caret Package. Journal of Statistical Software, 28(5), 1–26. https://doi.org/10.18637/jss.v028.i05

Li, J., Veeranampalayam-Sivakumar, A.-N., Bhatta, M., Garst, N. D., Stoll, H., Stephen Baenziger, P., Belamkar, V., Howard, R., Ge, Y., & Shi, Y. (2019). Principal variable selection to explain grain yield variation in winter wheat from features extracted from UAV imagery. Plant Methods, 15(1), 123. https://doi.org/10.1186/s13007-019-0508-7

Li, Q., Jin, S., Zang, J., Wang, X., Sun, Z., Li, Z., Xu, S., Ma, Q., Su, Y., Guo, Q., & Jiang, D. (2022). Deciphering the contributions of spectral and structural data to wheat yield estimation from proximal sensing. The Crop Journal, 10(5), 1334–1345. https://doi.org/https://doi.org/10.1016/j.cj.2022.06.005

Li, S., Yuan, F., Ata-UI-Karim, S. T., Zheng, H., Cheng, T., Liu, X., Tian, Y., Zhu, Y., Cao, W., & Cao, Q. (2019). Combining Color Indices and Textures of UAV-Based Digital Imagery for Rice LAI Estimation. Remote Sensing, 11(15), 1763. https://www.mdpi.com/2072-4292/11/15/1763

Liu, Y., Feng, H., Yue, J., Li, Z., Yang, G., Song, X., Yang, X., & Zhao, Y. (2022). Remote-sensing estimation of potato above-ground biomass based on spectral and spatial features extracted from high-definition digital camera images. Computers and Electronics in Agriculture, 198, 107089. https://doi.org/https://doi.org/10.1016/j.compag.2022.107089

Meier, U., Bleiholder, H., Buhr, L., Feller, C., Hack, H., Heß, M., Lancashire, P. D., Schnock, U., Stauß, R., & Van Den Boom, T. (2009). The BBCH system to coding the phenological growth stages of plants–history and publications. Journal für Kulturpflanzen, 61(2), 41–52.

Millet, E. J, Rodriguez Alvarez, M. X., Perez Valencia, D. M., Sanchez, I., Hilgert, N., van Rossum, B.-J., van Eeuwijk, F., & Boer, M. (2022). statgenHTP: High Throughput Phenotyping (HTP) Data Analysis. In https://biometris.github.io/statgenHTP/index.html, https://github.com/Biometris/statgenHTP/

Nguy-Robertson, A., Gitelson, A., Peng, Y., Viña, A., Arkebauer, T., & Rundquist, D. (2012). Green Leaf Area Index Estimation in Maize and Soybean: Combining Vegetation Indices to Achieve Maximal Sensitivity. Agronomy Journal, 104(5), 1336–1347. https://doi.org/https://doi.org/10.2134/agronj2012.0065

Pan, Y., Wu, W., Zhang, J., Zhao, Y., Zhang, J., Gu, Y., Yao, X., Cheng, T., Zhu, Y., Cao, W., & Tian, Y. (2023). Estimating leaf nitrogen and chlorophyll content in wheat by correcting canopy structure effect through multi-angular remote sensing. Computers and Electronics in Agriculture, 208, 107769. https://doi.org/https://doi.org/10.1016/j.compag.2023.107769

Pang, J., Palta, J. A., Rebetzke, G. J., & Milroy, S. P. (2014). Wheat genotypes with high early vigour accumulate more nitrogen and have higher photosynthetic nitrogen use efficiency during early growth. Functional Plant Biology, 41(2), 215–222. https://doi.org/https://doi.org/10.1071/FP13143

Pavuluri, K., Chim, B. K., Griffey, C. A., Reiter, M. S., Balota, M., & Thomason, W. E. (2015). Canopy spectral reflectance can predict grain nitrogen use efficiency in soft red winter wheat. Precision Agriculture, 16(4), 405–424. https://doi.org/10.1007/s11119-014-9385-2

Pinter, P. J., Jackson, R. D., Idso, S. B., & Reginato, R. J. (1981). Multidate spectral reflectance as predictors of yield in water stressed wheat and barley. International Journal of Remote Sensing, 2(1), 43–48. https://doi.org/10.1080/01431168108948339

Prey, L., Hanemann, A., Ramgraber, L., Seidl-Schulz, J., & Noack, P. O. (2022). UAV-Based Estimation of Grain Yield for Plant Breeding: Applied Strategies for Optimizing the Use of Sensors, Vegetation Indices, Growth Stages, and Machine Learning Algorithms. Remote Sensing, 14(24), 6345. https://www.mdpi.com/2072-4292/14/24/6345

Prey, L., Hu, Y., & Schmidhalter, U. (2020). High-Throughput Field Phenotyping Traits of Grain Yield Formation and Nitrogen Use Efficiency: Optimizing the Selection of Vegetation Indices and Growth Stages [Original Research]. Frontiers in Plant Science, 10. https://doi.org/10.3389/fpls.2019.01672

R Core Team. (2021). R: A Language and Environment for Statistical Computing. In R foundation for statistical computing. https://www.R-project.org/

Raun, W. R., Solie, J. B., Johnson, G. V., Stone, M. L., Lukina, E. V., Thomason, W. E., & Schepers, J. S. (2001). In-Season Prediction of Potential Grain Yield in Winter Wheat Using Canopy Reflectance. Agronomy Journal, 93(1), 131–138. https://doi.org/https://doi.org/10.2134/agronj2001.931131x

Ray, D. K., Mueller, N. D., West, P. C., & Foley, J. A. (2013). Yield Trends Are Insufficient to Double Global Crop Production by 2050. PLoS One, 8(6), e66428. https://doi.org/10.1371/journal.pone.0066428

Rischbeck, P., Elsayed, S., Mistele, B., Barmeier, G., Heil, K., & Schmidhalter, U. (2016). Data fusion of spectral, thermal and canopy height parameters for improved yield prediction of drought stressed spring barley. European Journal of Agronomy, 78, 44–59. https://doi.org/https://doi.org/10.1016/j.eja.2016.04.013

Satorre, E. H., & Slafer, G. A. (1999). Wheat: ecology and physiology of yield determination. CRC Press.

Shibayama, M., Salli, A., Häme, T., Iso-Iivari, L., Heino, S., Alanen, M., Morinaga, S., Inoue, Y., & Akiyama, T. (1999). Detecting Phenophases of Subarctic Shrub Canopies by Using Automated Reflectance Measurements. Remote Sensing of Environment, 67(2), 160–180. https://doi.org/https://doi.org/10.1016/S0034-4257(98)00082-0

Spano, G., Di Fonzo, N., Perrotta, C., Platani, C., Ronga, G., Lawlor, D. W., Napier, J. A., & Shewry, P. R. (2003). Physiological characterization of ‘stay green’ mutants in durum wheat. Journal of Experimental Botany, 54(386), 1415–1420. https://doi.org/10.1093/jxb/erg150

Taniguchi, S., Sakamoto, T., Imase, R., Nonoue, Y., Tsunematsu, H., Goto, A., Matsushita, K., Ohmori, S., Maeda, H., Takeuchi, Y., Ishii, T., Yonemaru, J.-i., & Ogawa, D. (2022). Prediction of heading date, culm length, and biomass from canopy-height-related parameters derived from time-series UAV observations of rice [Original Research]. Frontiers in Plant Science, 13. https://doi.org/10.3389/fpls.2022.998803

Tucker, C. J. (1979). Red and photographic infrared linear combinations for monitoring vegetation. Remote Sensing of Environment, 8(2), 127–150. https://doi.org/https://doi.org/10.1016/0034-4257(79)90013-0

Vatter, T., Gracia-Romero, A., Kefauver, S. C., Nieto-Taladriz, M. T., Aparicio, N., & Araus, J. L. (2022). Preharvest phenotypic prediction of grain quality and yield of durum wheat using multispectral imaging. Plant J, 109(6), 1507–1518. https://doi.org/10.1111/tpj.15648

Walter, A., Liebisch, F., & Hund, A. (2015). Plant phenotyping: from bean weighing to image analysis. Plant Methods, 11(1), 14. https://doi.org/10.1186/s13007-015-0056-8

Wang, F., Li, W., Liu, Y., Qin, W., Ma, L., Zhang, Y., Sun, Z., Wang, Z., Li, F., & Yu, K. (2022). Characterization of N distribution in different organs of winter wheat using UAV-based remote sensing. bioRxiv, 2022.2011.2002.514839. https://doi.org/10.1101/2022.11.02.514839

Wang, F., Yi, Q., Hu, J., Xie, L., Yao, X., Xu, T., & Zheng, J. (2021). Combining spectral and textural information in UAV hyperspectral images to estimate rice grain yield. International Journal of Applied Earth Observation and Geoinformation, 102, 102397. https://doi.org/https://doi.org/10.1016/j.jag.2021.102397

Watt, M., Fiorani, F., Usadel, B., Rascher, U., Muller, O., & Schurr, U. (2020). Phenotyping: New Windows into the Plant for Breeders. Annual Review of Plant Biology, 71(1), 689–712. https://doi.org/10.1146/annurev-arplant-042916-041124

Wu, C., Niu, Z., Tang, Q., & Huang, W. (2008). Estimating chlorophyll content from hyperspectral vegetation indices: Modeling and validation. Agricultural and forest meteorology, 148(8), 1230–1241. https://doi.org/https://doi.org/10.1016/j.agrformet.2008.03.005

Yue, J., Yang, G., Tian, Q., Feng, H., Xu, K., & Zhou, C. (2019). Estimate of winter-wheat aboveground biomass based on UAV ultrahigh-ground-resolution image textures and vegetation indices. ISPRS Journal of Photogrammetry and Remote Sensing, 150, 226–244. https://doi.org/https://doi.org/10.1016/j.isprsjprs.2019.02.022

Zhang, J., Qiu, X., Wu, Y., Zhu, Y., Cao, Q., Liu, X., & Cao, W. (2021). Combining texture, color, and vegetation indices from fixed-wing UAS imagery to estimate wheat growth parameters using multivariate regression methods. Computers and Electronics in Agriculture, 185, 106138. https://doi.org/https://doi.org/10.1016/j.compag.2021.106138

Zheng, H., Cheng, T., Zhou, M., Li, D., Yao, X., Tian, Y., Cao, W., & Zhu, Y. (2019). Improved estimation of rice aboveground biomass combining textural and spectral analysis of UAV imagery. Precision Agriculture, 20(3), 611–629. https://doi.org/10.1007/s11119-018-9600-7

Zhou, X., Kono, Y., Win, A., Matsui, T., & Tanaka, T. S. T. (2021). Predicting within-field variability in grain yield and protein content of winter wheat using UAV-based multispectral imagery and machine learning approaches. Plant Production Science, 24(2), 137–151. https://doi.org/10.1080/1343943X.2020.1819165

